# Ecology and Epidemiology of Wheat Curl Mite and Mite-Transmissible Viruses in Colorado and Insights into the Wheat Virome

**DOI:** 10.1101/2020.08.10.244806

**Authors:** Tessa Albrecht, Samantha White, Marylee Layton, Mark Stenglein, Scott Haley, Punya Nachappa

**Author notes:** Corresponding author: Punya Nachappa. Funding: Colorado Wheat Research Foundation, Colorado Wheat Administrative Committee.

## Abstract

The wheat curl mite (WCM)-transmissible wheat streak disease complex is the most serious disease of wheat in the U.S. Great Plains. In the current study, we determined the genetic variability in WCM and mite-transmitted viruses in Colorado and identified sources of resistance in Colorado wheat germplasm to wheat streak disease complex. We identified two distinct genotypes of WCM, Type 1 and Type 2 based on the ribosomal ITS1 region. Both genotypes were found to co-exist throughout the wheat producing regions of Colorado. Analysis of the whole genome and partial coat protein sequences revealed rich diversity of wheat streak mosaic virus (WSMV) and High Plains wheat mosaic virus (HPWMoV) isolates collected from Colorado, whereas triticum mosaic virus (TriMV) showed low sequence variability. Analysis of WSMV isolates revealed two novel isolates and one that was 100% similar to a new variant of WSMV from Kansas. Interestingly, between 2-4 genotypes of all 8 RNA segments of HPWMoV were identified, which suggests new variants of emaraviruses and co-occurrence of multiple strains within host populations. Several novel viruses including mycoviruses were identified for the first time in Colorado. We found variation in WSMV resistance among wheat varieties; however a variety that harbored dual resistance to mite and WSMV had lower virus titer compared to varieties that contained single resistance gene. This suggests that pyramiding genes will ensure improved and durable resistance. Future research may be aimed at elucidating the dynamics, diversity, and distribution of the new WSMV and HPWMoV isolates and their responses to wheat genotypes.

---

Wheat (*Triticum aestivum* L.) is considered the most important crop in the 21^st^ century as it serves as a nutritional source of calories and protein in the human diet worldwide (Arzani and Ashraf 2017; Curtis and Halford 2014). In the United States, wheat ranks third among field crops in planted acreage, production, and gross farm receipts, behind corn and soybeans (USDA-ERS 2019). Among the top 10 wheat growing states, Colorado ranked 6^th^ in 2019 with 2,150,000 acres being planted and a yield of 49 bushels per acre resulting in total production of 98,000,000 bushels valued at $387,100,000 (USDA-NASS 2019). The wheat curl mite (WCM), *Aceria tosichella* Keifer (Acari: Eriophyidae) is a globally important pest affecting wheat production in the Americas, Europe, and Asia (Skoracka et al. 2018). The mite causes direct damage by feeding, which can reduce cereal yield (Harvey et al. 2000). But more importantly, WCM-transmitted viruses including wheat streak mosaic virus (family Potyviridae/genus Tritimovirus; acronym WSMV) (Slykhuis 1955), triticum mosaic virus (Potyviridae/Poacevirus; TriMV) (Seifers et al. 2009) and High plains wheat mosaic virus (Fimoviridae/Emaravirus; HPWMoV) (Seifers et al. 1997) are among the most significant viruses in U.S. agriculture, responsible for yield losses in wheat, barley, oats and rye (Burrows et al. 2009; Navia et al. 2013). Average yield losses from the WCM-WSMV complex range from 5 to 7% in the US Great Plains, but 100% yield losses may occur in some fields (Appel et al. 2015).

Worldwide, the WCM has been found to be a diverse species complex with numerous genetic lineages (Skoracka et al. 2018). In North America however, only two genetically distinct genotypes of WCM have been characterized based on ribosomal ITS1 and mitochondrial Cytochrome oxidase I/II partial sequences: Type 1, initially identified from South Dakota, Kansas, Montana, Nebraska and Texas, and Type 2, from Nebraska (Hein et al. 2012). Both genotypes occur in mixed populations in wheat-producing areas of the U.S. Great Plains. The two distinct genotypes demonstrate different responses to curl mite colonization (*Cmc*) genes; *Cmc1, Cmc2, Cmc3* and *Cmc4* (Dhakal et al. 2017; Harvey et al. 1999) and differential viral transmission efficiencies (Hein et al. 2012; McMechan et al. 2014; Seifers et al. 2002; Wosula et al. 2016). For example, Type 2 is more virulent and makes wheat lines carrying the 1AL.1RS (*Cmc3* resistance gene) susceptible (Dhakal *et al*., 2017) and Type 2 mites transmit WSMV at higher rates compared to Type 1 mites (Wosula *et al*., 2016).

The WSMV populations are complex as well with numerous genotypes (Robinson and Murray 2013; Schubert et al. 2015), although different genotypes rarely occur in the same plant (McNeil et al. 1996). In the U.S., there are two WSMV isolates, Sidney 81 and Type, sharing 97.6% nucleotide sequence identity, and produce similar symptoms in wheat (Choi et al. 2001; Hall et al. 2001). A third isolate, El Batán, from Mexico has diverged from the American strains and has 79% nucleotide sequence identity to Sidney 81 and Type (Choi et al. 2001). In contrast, TriMV field populations showed minimal amounts of sequence variation suggesting that the populations are very homogenous (Fuentes-Bueno et al. 2011). There is little information about the phylogenetic relationships between HPWMoV isolates. There appears to be two distinct groups of HPWMoV isolates within the U.S. (Stewart 2016). Currently there are three sources of host resistance to WSMV - *Wsm1, Wsm2*, and *Wsm3* (Liu et al. 2011; Lu et al. 2011; Triebe et al. 1991). However, some of these resistance alleles are temperature sensitive and do not prevent virus infection and replication above 18°C (Fahim et al. 2012). More recently, a novel QTL was identified on wheat chromosome 6DS from the wheat cultivar, TAM112, which provides WCM resistance and moderate WSMV resistance (Dhakal et al. 2018). Genes for resistance to TriMV and HPWMoV have not been identified.

One of the most effective ways of controlling WCM-virus complex is by planting mite and disease resistant varieties; however, knowledge of mite and virus genotypes occurring in a given area is critical because these genetic differences correspond to biological responses at the phenotypic level (Hein et al. 2012). While Colorado is a major wheat producing state, there is no information about the WCM-virus complex in the region. Moreover, little is known about emerging and/or novel viruses of wheat in Colorado. Next generation sequencing is a powerful tool that allows researchers to detect and characterize novel viruses (and bacterial and fungal pathogens) and explore their diversity and pathogenicity in agricultural crops (Villamor et al. 2019). NGS is finding increased applications in revealing the viromes that contribute to the disease phenotype. The term “virome” is defined as the genomes of all the viruses inhabiting a specific organism or environment. In the current study, we determined the genetic variability in WCM and mite-transmitted viruses in Colorado and identified sources of resistance in Colorado wheat germplasm to WSMV and TriMV. In addition, we investigated the viromes of wheat from four different locations in Colorado. To our knowledge, our study is among the first to report on the wheat virome in the U.S.

## Materials and Methods

### Wheat Curl Mite and Plant Tissue Collection

Symptomatic wheat leaf tissues were collected across eastern Colorado by researchers, extension agents and producers and delivered to our laboratory at Colorado State University. Plants were examined using a dissecting microscope for the presence of WCMs. If present, mites were transferred to healthy wheat plants of susceptible wheat varieties, Pronghorn or Hatcher at the four-leaf stage or older. Plants were grown in gallon pots with 2-3 plants per pot in Promix HP^©^ soil. A minimum of 10 mites were transferred either singly using a fine camel hairbrush or by placing a small section of the WCM-infested wheat leaf onto healthy host plants. Mite colonies were maintained on wheat plants in 45.72 cm x 45.72 cm x 76.20 cm insect cages with no-thrips insect screen (Bioquip, CA, USA) under 16:8 (Light: Dark) hour (h) cycle at approximately 23°C under laboratory conditions. Additionally, leaf tissues were tested for presence of WSMV, TriMV and HPWMoV. Approximately 40 mg of the most symptomatic leaf tissue was collected for RNA extraction and virus detection.

### Mite and Virus Genotyping and Phylogenetic Analysis

Wheat curl mite DNA was extracted from single mites using the MyTaq™ Extract-PCR Kit (Bioline Meridian Bioscience, London, UK)) according to the manufacturer’s recommendations with the exception of the amount of starting material being less than 3 mg. The ribosomal ITS1 partial sequences were amplified from mite DNA using the primers listed in Table 1. To identify virus isolates present in the wheat samples, the NIb (Nuclear Inclusion putative polymerase) region of WSMV and partial sequences of TriMV coat protein (CP) and HPWMoV nucleoprotein (NP) were amplified. The genotype each of the WCMs and associated viruses from the field samples were determined by sequencing the resulting amplicons (GeneWiz, NJ, USA). Sequence alignments were generated using ClustalW in MEGA X (Kumar et al. 2018) followed by phylogenetic analyses based on the Maximum Likelihood method with 1000 replications.

**Table 1.**
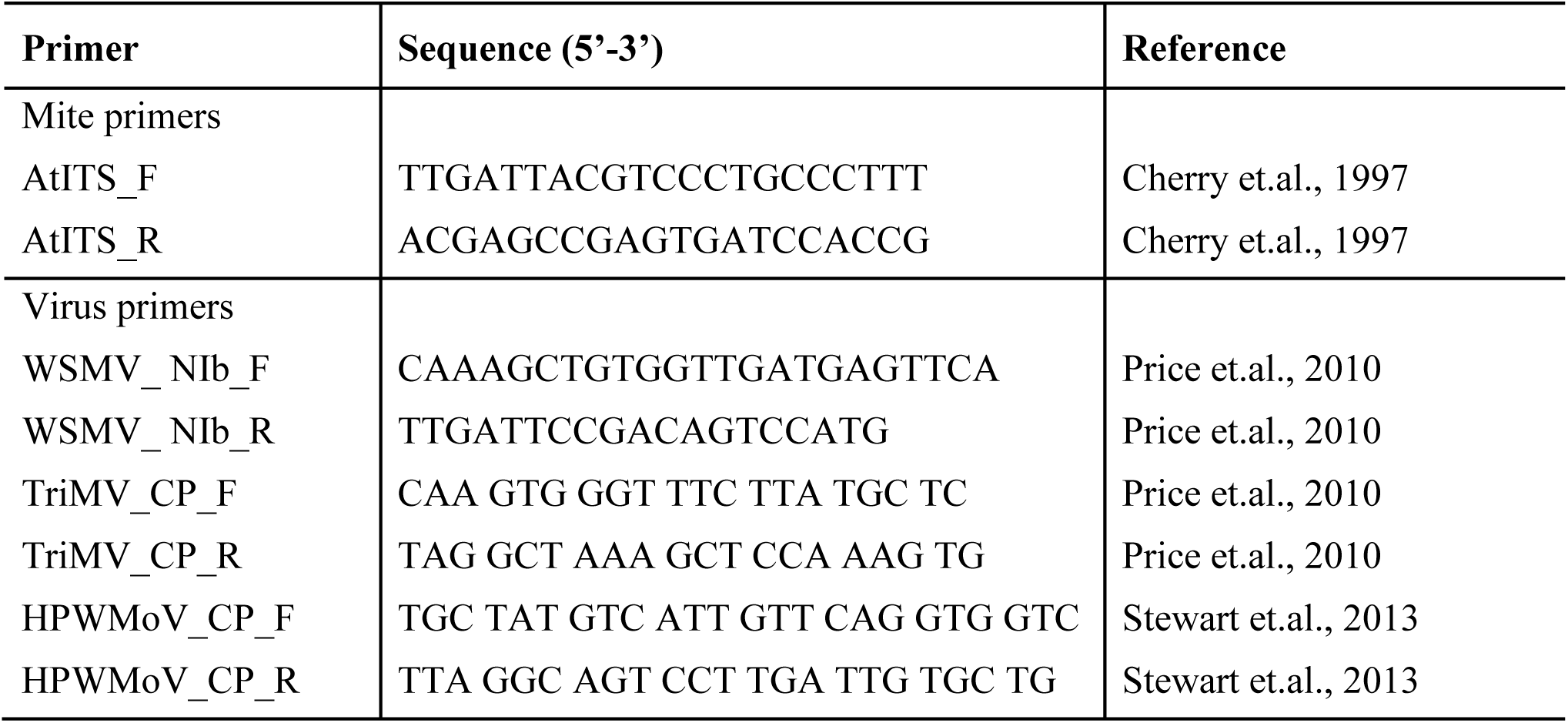
Primers used for PCR analysis in this study.

### Virus Detection and Quantification

Total RNA was extracted from approximately 40 mg homogenized leaf tissues obtained from various sources described above, lysed in Trisure® (Bioline Meridian Bioscience) using Direct-zol® RNA Purification Kit (Zymo Research, CA, USA) according to the manufacturer’s recommendations. The quantity of RNA was approximated using a NanoDrop One spectrophotometer (Thermo Fisher Scientific, MA, USA) and stored at −80°C until virus quantification. To detect and quantify WSMV and TriMV, approximately 50 ng of RNA was used in a previously published qRT-PCR duplex assay (Price et al. 2010) with the TaqMan® RNA-to-Ct™ 1-step kit (Applied Biosystems™, ThermoFisher Scientific) on a QuantStudio™ 3 Real-Time PCR system (Applied Biosystems™, ThermoFisher Scientific). Reaction condition were set to incubate for the RT reaction at 48°C for 30 min, initial denaturation at 95°C for 10 min, and 40 cycles of denaturation at 95°C for 15 s and anneal/extension at 60°C for 1 min. Field samples with C_Q_ values above the lower detection limit, as defined by the standard curves described below, were considered to be positive for the specified virus. To quantify virus titer of WSMV and TriMV in samples, a standard curve was generated using a 319 bp amplicon of the NIb region of WSMV and a 677 bp amplicon of the CP of TriMV, both containing the respective qPCR target. Primers used to produce each amplicon are listed in Table 1. Ten-fold serial dilutions of each target amplicon ranging from 500 fg to 0.05 fg of DNA was used to generate a standard curve relating C_Q_ values to the estimated copy number corresponding to each concentration of target DNA as per Keough et al. (2016).

To detect HPWMoV, complementary DNA (cDNA) was synthesized from 1 μg of total RNA using the Verso® cDNA Synthesis Kit (ThermoFisher Scientific). The partial sequence encoding the nucleoprotein from HPWMoV was amplified using specific primers (Table 1) from two-fold diluted cDNA with GoTaq® Flexi DNA polymerase (Promega, WI, USA) with the following reaction conditions: initial denaturation at 95°C for two minutes, followed by 35 cycles of denaturation at 95°C for 30 seconds, annealing at 55°C for 30 seconds and extension at 72°C for one minute, and final extension at 72°C for five minutes. Products were visualized on a 1% agarose gel with positive and negative controls to determine the presence of HPWMoV in each sample.

### Wheat Virome Analysis

Four leaf tissue samples that previously tested positive for WSMV from Larimer county., positive for WSMV and TriMV from Bent county., positive for WSMV and HPWMoV from Phillips county., and positive for WSMV and TriMV from Kit Carson county. were used for wheat virome analysis. Total RNA was extracted as described above and checked for quality using a Nanodrop One spectrophotometer (ThermoFisher Scientific) and quantity using a Qubit fluorometer (ThermoFisher Scientific). Approximately 2 µg of RNA was submitted to the CSU Next Generation Sequencing Facility, where library preparation, quality measurements, and sequencing was performed. Briefly, RNA quality was confirmed using an Aligent Tapestation instrument. Shotgun RNA libraries were constructed using the Kapa Biosystems RNA HyperPrep kit (Roche, IN, USA) according to the manufacturer’s instructions. Pooled libraries were sequenced on an Illumina NextSeq 500 instrument to produce single-end 150 nucleotide (nt) reads. Datasets contained an average of 9.4×10^6^ reads.

### Bioinformatic Analyses

Virus and virus-like sequences were identified as previously described (Cross et al. 2018). Analysis scripts are available at https://github.com/stenglein-lab/taxonomy_pipeline/. Low quality and adapter sequences were removed using cutadapt software (Martin 2011), leaving an average of 8.3×10^6^ sequences remaining per dataset (93%). Duplicate reads were collapsed with cd-hit (Li and Godzik 2006), leaving an average of 1.8×10^6^ unique reads per dataset (20%). Host (wheat)-derived reads were removed by bowtie2 alignment (Langmead and Salzberg 2012) to the *Triticum aestivum* reference genome (assembly accession GCA_900519105.1) (Appels et al. 2018). After all filtering operations, an average of 0.25×10^6^ reads (3%) remained per dataset. Remaining non-host reads were assembled into contigs using the Spades assembler (Bankevich et al. 2012). Contigs and non-assembling reads were taxonomically categorized first by nucleotide-level alignment to the NCBI nucleotide (nt) database using BLASTN, and then by protein-level alignment to the NCBI protein (nr) database using the diamond aligner (Altschul et al. 1990; Buchfink et al. 2015). This produced a comprehensive metagenomic classification of all non-host reads. Although we focused on viruses, this also constitute a valuable dataset about the entire wheat-associated microbiota (bacteria, fungi, etc.) for future use by us and others. Candidate virus sequences were manually validated by aligning reads to draft genome assemblies using bowtie2. Phylogenetic trees were constructed to reveal the relationships of identified viruses with other known isolates using the ClustalW software program. Then, analysis of SNPs for the viruses from assembled virome data were performed using the Tablet software program to determine genetic diversity. Lastly, the raw sequence data was deposited in the NCBI Sequence Read Archive (SRA) repository under submission number SUB7870854. The annotated viromes obtained from the study were deposited in GenBank at NCBI with respective accession numbers MT762109-MT762125 and MT822723-MT822732 (awaiting three additional accession numbers).

### Wheat Germplasm and Virulence Test

A natural infection of wheat streak mosaic virus was observed in the Colorado State University Irrigated Variety Performance Trial (IVPT) at Burlington, CO, in 2019. The trial included 24 different genotypes (released varieties and experimental lines), planted in a randomized complete block design with three replications. Each plot was 7 rows wide, 10.7 m long, with an inter-row row spacing of 0.23 m. The trial was planted on October 3, 2018, at a seeding rate of approximately 2.9 million seeds per hectare. Symptoms of infection of wheat streak mosaic virus were first observed on May 15, 2019, shortly after the heading growth stage (Zadoks 50-60), and visual observations of symptom expression were recorded on June 14, 2019. Symptom expression was recorded on a 1 (most resistant)-9 (most susceptible) scale based on yellow streaking or mosaic patterns on the leaves, variable plant height within each plot (stunting), and tillering.

To quantify virus titer in wheat varieties, 10 leaf samples were collected on June 21, 2019. Samples were collected without regard to visual disease symptoms in a diagonal line stretching between opposite corners of each plot. Tissue was collected from all leaves representing a single plot for RNA extraction and virus quantification. All ten leaves were stacked, and a small section of tissue was cut from the center of the stack. Total RNA extraction and virus quantification was conducted as described above. The difference in log copy number of viral RNA among wheat lines was analyzed using two-way ANOVA (PROC GLM) with line/variety and plots/replicate as fixed effects and the interaction term. Treatment comparisons were performed using Tukey’s family error rate (*P* < 0.05).

## Results

### Identification of Mite Genotypes

To explore the genotypic variability in our regional mite populations, we analyzed the ribosomal ITS1 regions from six WCM populations collected at various field locations including, Larimer, Kit Carson, Adams, Phillips, and Sedgwick counties throughout Colorado. Phylogenetic relationships among ITS1 sequences for Colorado populations (accessions MT465683-MT465687) and representative sequences from different states in the U.S. and around the world are shown in Figure 1. Overall, there was limited variability among the ITS1 sequences, but there is a clear distinction between Type 1 and Type 2 mites. The WCM populations collected from two counties, Larimer and Sedgwick, had the same ITS1 sequence as that of Type 1 mites from Texas, Nebraska, Kansas, Montana and South Dakota. In contrast, mites collected from four Colorado counties, Larimer, Kit Carson, Adams, and Phillips had ITS1 sequences identical to that of Type 2 mites from Nebraska (Fig. 1). Mites collected from Larimer Co. during different times of the growing season belonged to both genotypes suggesting that both Type 1 and Type 2 genotypes co-exist throughout the wheat producing regions of Colorado similar to the other wheat producing regions of the U.S. Great Plains.

**Fig. 1.**
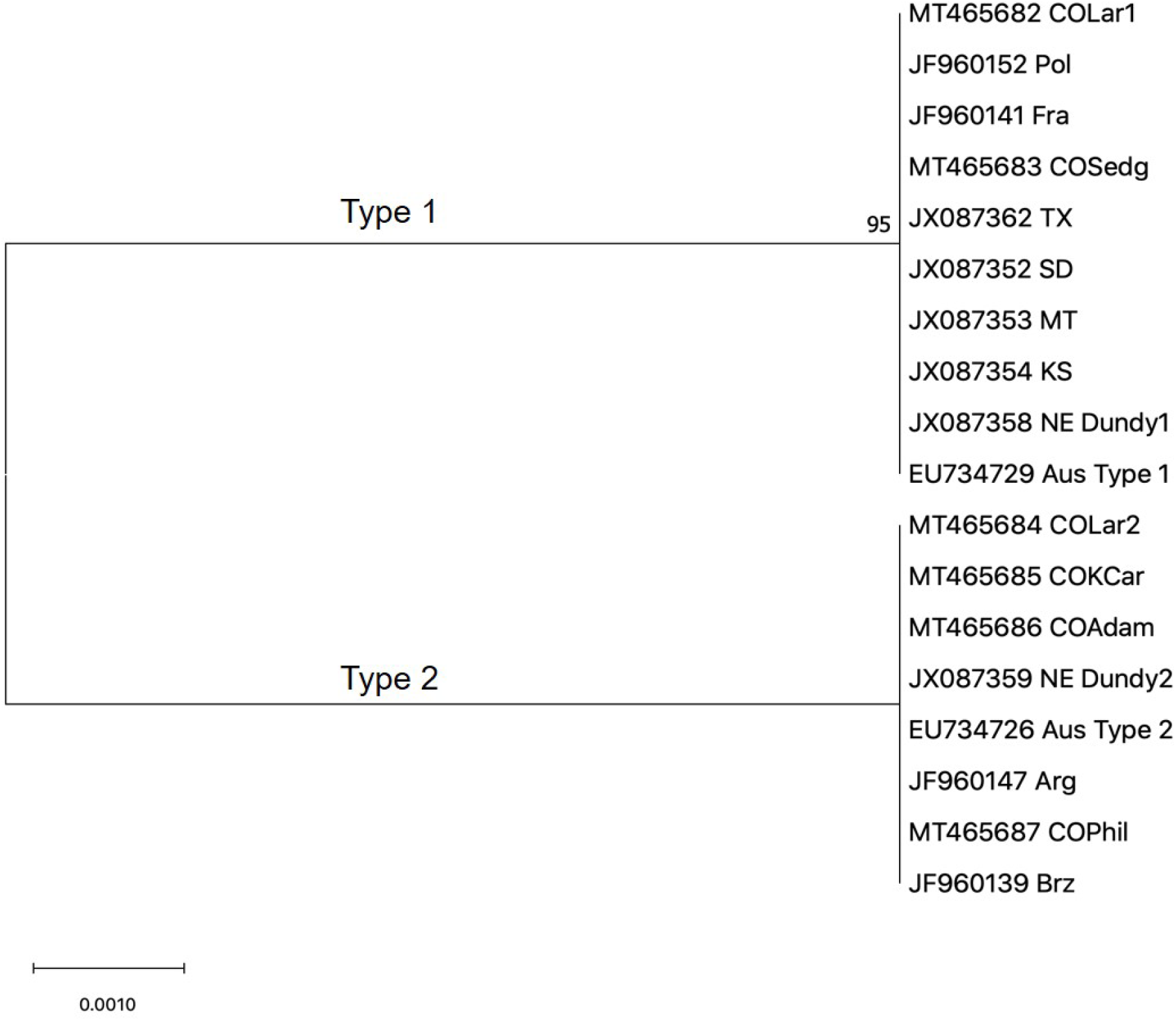
Phylogenetic tree of wheat curl mite populations using the ribosomal internal transcribed spacer (ITS1) region. Scale bar indicates percent genetic distances. Phylogenetic analysis by maximum likelihood method was based on a sequence alignment using ClustalW in MEGAX. Bootstrap values less than 70% out of 1000 replicates are not shown.

### Virus Occurence in Colorado

Survey of WCM-transmitted viruses revealed the presence of all three economically-important viruses in Colorado, WSMV, TriMV and HPWMoV (Fig. 2). Of the 40 symptomatic samples tested, 38 were positive for one or more WCM-transmitted viruses. WSMV was found in all surveyed counties (Fig. 2). Coinfection of WSMV and TriMV was detected in seven samples from Bent, Kiowa, Kit Carson and Weld counties. Only three samples were positive for both WSMV and HPWMoV from the Phillips county indicating low incidence of coinfection of these viruses. No single infection or coinfection of TriMV and HPWMoV was detected. We made an intriguing observation in that samples collected early in the season (April 2018 and May 2018) were infected with only WSMV, while samples collected later in the season (past June 2018) were positive for coinfection of WSMV and TriMV.

**Fig. 2.**
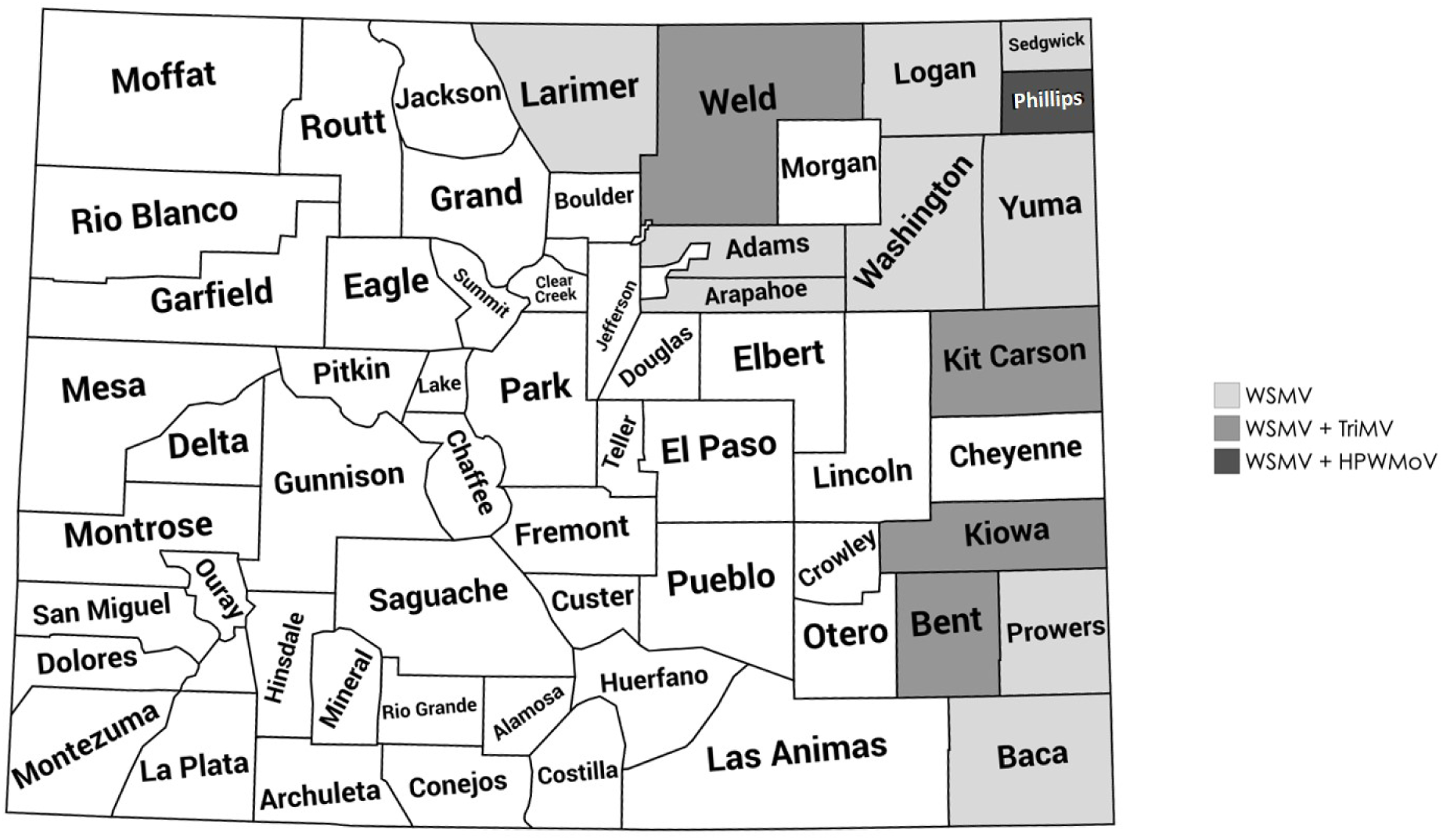
Map of occurrence of wheat curl mite-transmitted viruses in eastern Colorado during 2019 as determined by RT-qPCR analysis Symptomatic leaf tissues were collected from various locations or obtained from extension agents and growers. The number of samples tested in each county ranged from one to six. Map was generated using mapchart.net.

### Identification of Virus Genotypes

To determine the genetic variability among WSMV isolates in Colorado, we sequenced a portion of the WSMV-NIb region and performed phylogenetic analyses with isolates from other wheat producing states in the U.S. and other regions of the world. Phylogenetic analysis revealed diversity among Colorado isolates from Larimer, Sedgwick, Kiowa, Kit Carson, and Phillips counties; (MT465688-MT465692) and other available sequences in GenBank (Fig. 3). Interestingly, an isolate, Larimer county 1, was 100% similar to a Kansas isolate (MK318278) that was collected from a wheat variety carrying the *Wsm2* virus resistance gene that is known to confer resistance to WSMV (Fellers et al. 2019). Another isolate from Kit Carson county with similarity to isolates from neighboring states was collected from Snowmass 2.0, another variety carrying the *Wsm2* virus resistance gene. The isolate Larimer county 2 collected from same location as the Larimer county 1 isolate, appeared to be genetically distinct from other isolates in the U.S. The Phillips county isolate is also genetically distinct from the others (Fig. 3).

**Fig. 3.**
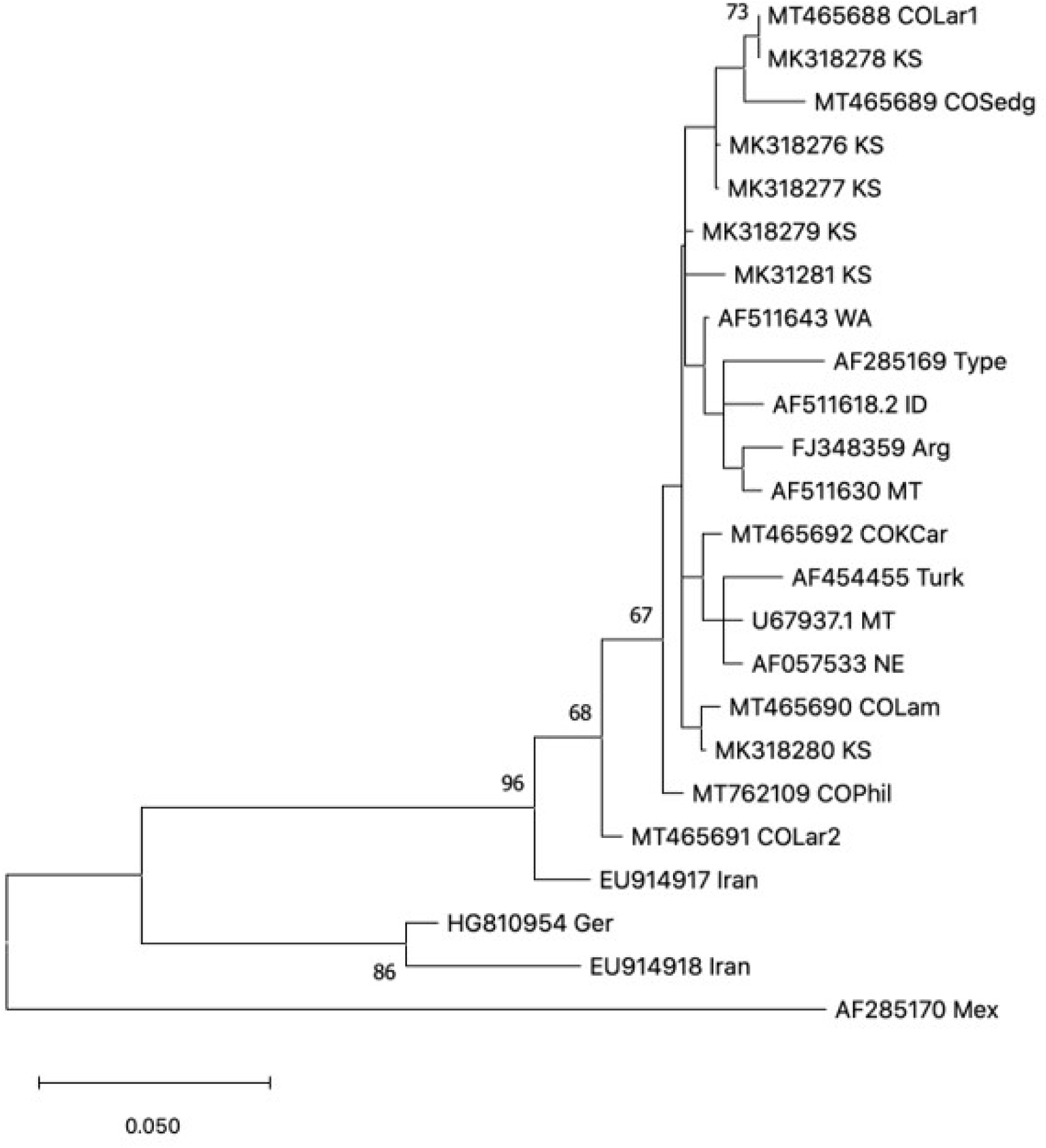
Phylogenetic tree of wheat streak mosaic virus (WSMV) isolates using WSMV-NIb region. Scale bar indicates percent genetic distances. Phylogenetic analysis by maximum likelihood method was based on a sequence alignment using ClustalW in MEGAX. Bootstrap values less than 65% out of 1000 replicates are not shown.

In contrast, to the genetic diversity in WSMV isolates, there was limited variability in TriMV isolates collected in Colorado compared to that of other sequenced isolates. There was high sequence similarity between the TriMV isolate from Kit Carson county (MT563401) and the other available TriMV sequences from surrounding states, with a range of 99.44% to 98.68% identity in a 531bp region of the coat protein.

Phylogenetic analysis reveals two distinct groups of the HPWMoV isolates among available HPWMoV nucleoprotein sequences as observed by previous studies (Stewart 2016). Analysis of the HPWMoV NP sequence obtained from Phillips county (MT563400) revealed high identity (98.97-98.29%) to isolates in the group from Ohio and Texas, and only 83.28% identity to isolates in the other group from Kansas and Nebraska as well as Ohio (Data not shown). Phylogenetic analysis using the complete HPWMoV RNA3 segment encoding the nucleoprotein is described in detail below.

### Wheat Virome Analysis

Wheat viromes were analyzed in four samples collected from Larimer, Bent, Phillips and Kit Carson counties. Table 2 summarizes the metatranscriptomic data descriptive statistics. NGS analysis revealed that the predominant viruses were WSMV and TriMV. WSMV was detected in all four locations (the Bent county sample had low WSMV titer and coverage was not sufficient for coding sequences to be assembled) and TriMV was detected in two out of four locations (Table 3). HPWMoV was only detected in Phillips county. Interestingly, this sample also contained several mycoviruses or fungus-infecting viruses including, Plasmopara viticola associated mononega virus 1-like (MT822729), Plasmopara viticola associated mitovirus 7 (MT822730), Plasmopara viticola associated ourmia-like virus, Fusarium poae negative-stranded virus 2-like (MT822731) and Coniothyrium diplodiella negative-stranded RNA virus (MT822732). These likely correspond to viruses infecting a *Fusarium* spp., which was present in this dataset as the most abundant non-host taxon identified. A new variant of Ixeridium yellow mottle virus 2-like was also identified in the Phillips sample, which had 45% amino acid identity to an unclassified umbravirus, ixeridium yellow mottle virus 2 (YP_009352229.1) (Table 3).

**Table 2.** Wheat shotgun metagenomic sequencing data quality.

| **Library/**<br>**county** | **Number of raw**<br>**reads** | **Number of reads**<br>**after filtering**<br>**low quality and**<br>**adapter**<br>**sequences** | **Number of**<br>**unique**<br>**sequences** | **Number of host**<br>**filtered**<br>**sequences** |
| --- | --- | --- | --- | --- |
| Bent | 8148234 | 7559238 | 2392705 | 12683 |
| Kit Carson | 9336195 | 8729024 | 1848876 | 214637 |
| Larimer | 9355272 | 8695140 | 1648474 | 14797 |
| Phillips | 10668454 | 10085067 | 1443655 | 744157 |

**Table 3.**
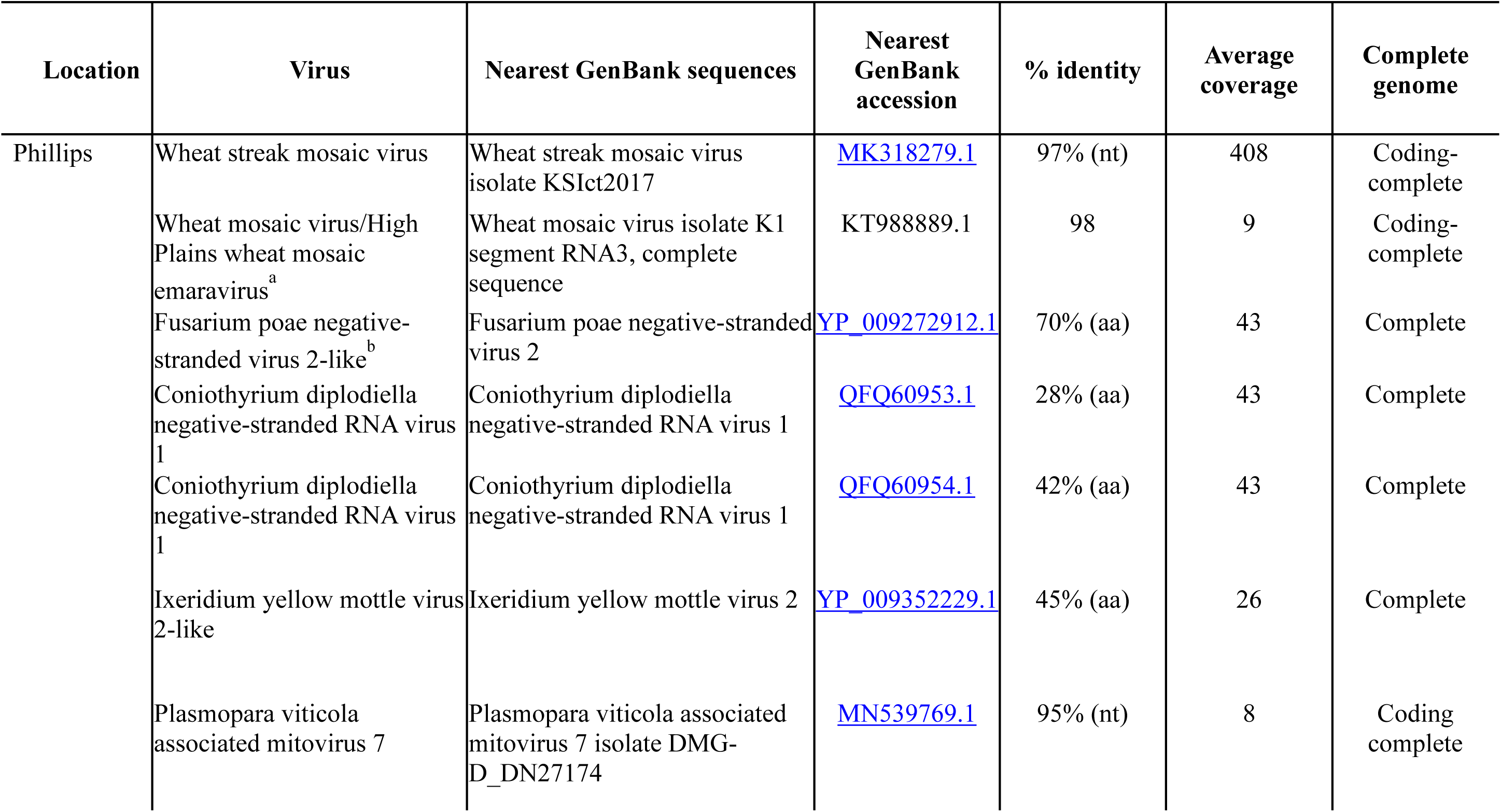
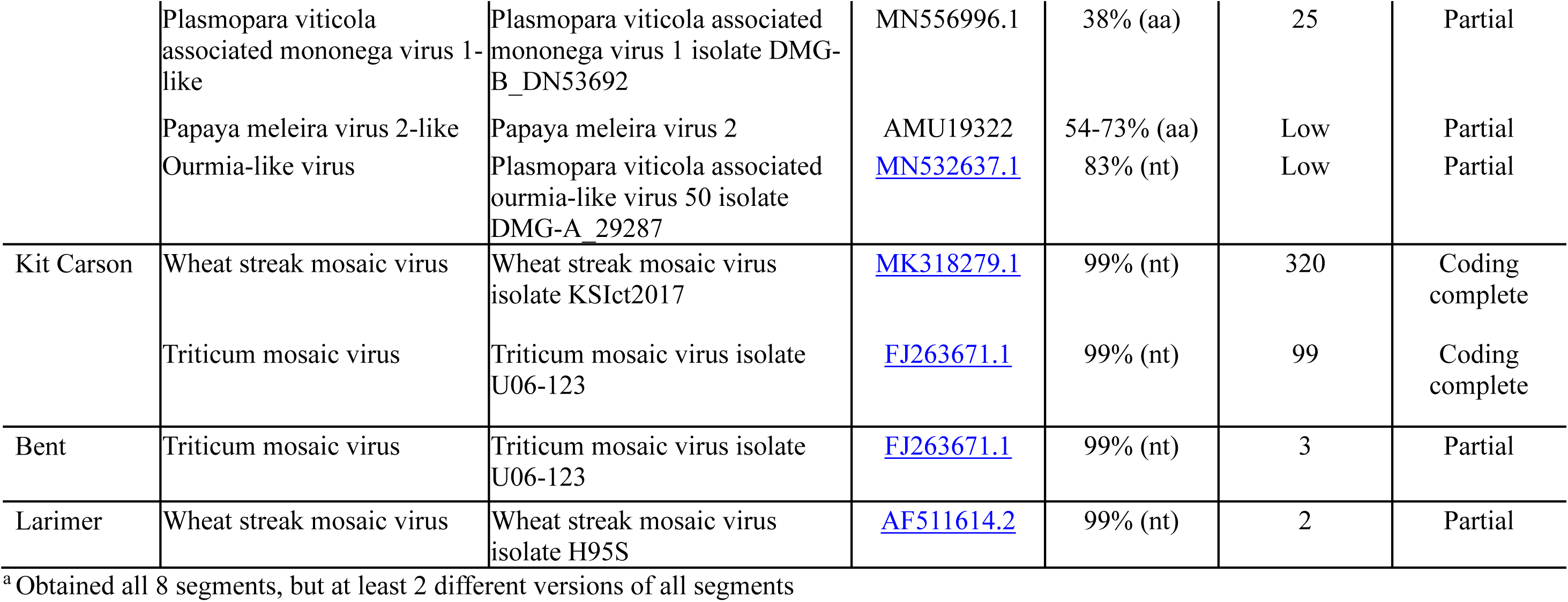
Summary of wheat viromes from Colorado.

| Location | Virus | Nearest GenBank sequences | Nearest GenBank accession | % identity | Average coverage | Complete genome |
| --- | --- | --- | --- | --- | --- | --- |
| Phillips | Wheat streak mosaic virus | Wheat streak mosaic virus isolate KS1ct2017 | MK318279.1 | 97% (nt) | 408 | Coding-complete |
|  | Wheat mosaic virus/High Plains wheat mosaic emaravirus<sup>a</sup> | Wheat mosaic virus isolate K1 segment RNA3, complete sequence | KT988889.1 | 98 | 9 | Coding-complete |
|  | Fusarium poae negative-stranded virus 2-like<sup>b</sup> | Fusarium poae negative-stranded virus 2 | YP_009272912.1 | 70% (aa) | 43 | Complete |
|  | Coniothyrium diplodiella negative-stranded RNA virus 1 | Coniothyrium diplodiella negative-stranded RNA virus 1 | QFQ60953.1 | 28% (aa) | 43 | Complete |
|  | Coniothyrium diplodiella negative-stranded RNA virus 1 | Coniothyrium diplodiella negative-stranded RNA virus 1 | QFQ60954.1 | 42% (aa) | 43 | Complete |
|  | Ixeridium yellow mottle virus 2-like | Ixeridium yellow mottle virus 2 | YP_009352229.1 | 45% (aa) | 26 | Complete |
|  | Plasmopara viticola associated mitovirus 7 | Plasmopara viticola associated mitovirus 7 isolate DMG-D_DN27174 | MN539769.1 | 95% (nt) | 8 | Coding complete |
|  | Plasmopara viticola associated mononega virus 1-like | Plasmopara viticola associated mononega virus 1 isolate DMG-B_DN53692 | MN556996.1 | 38% (aa) | 25 | Partial |
|  | Papaya meleira virus 2-like | Papaya meleira virus 2 | AMU19322 | 54-73% (aa) | Low | Partial |
|  | Ourmia-like virus | Plasmopara viticola associated ourmia-like virus 50 isolate DMG-A_29287 | MN532637.1 | 83% (nt) | Low | Partial |
| Kit Carson | Wheat streak mosaic virus | Wheat streak mosaic virus isolate KS1ct2017 | MK318279.1 | 99% (nt) | 320 | Coding complete |
|  | Triticum mosaic virus | Triticum mosaic virus isolate U06-123 | FJ263671.1 | 99% (nt) | 99 | Coding complete |
| Bent | Triticum mosaic virus | Triticum mosaic virus isolate U06-123 | FJ263671.1 | 99% (nt) | 3 | Partial |
| Larimer | Wheat streak mosaic virus | Wheat streak mosaic virus isolate H95S | AF511614.2 | 99% (nt) | 2 | Partial |

<sup>a</sup> Obtained all 8 segments, but at least 2 different versions of all segments

We were able to assemble the complete or near complete genome of several of the viruses using NGS data. We obtained complete coding sequences for the polyprotein of WSMV from two of the four samples that were submitted, Kit Carson and Phillips counties (MT762110 and MT762109). Partial sequences for much of the polyprotein for the Larimer isolate were also assembled (MT822723-MT822728). We aligned the Colorado WSMV sequences with those from Kansas (Fellers et.al. 2019), and WSMV reference strains, Type (AF285169) and Sidney 81 (AF057533). A common amino acid change at position 2235 reported by Fellers et al.(2019), between Sidney 81 and other isolates was also present in translated amino acid alignment of our Colorado isolates (Fig. 4). Where Sidney 81 contains threonine (T), other isolates have either valine (V) or methionine (M). Notably, the Phillips isolate contained 18 unique amino acid changes throughout the polyprotein of the twelve aligned sequences, primarily in the P1 protein and HC-Pro regions. These findings are in agreement with alignment of WSMV-NIb sequences which also showed simialrities between Colorado isolate and a new variant of WSMV detected in Kansas. It also confirms that the WSMV isolate from Phillips county is highly divergent (Fig. 3). The biological significance of genotypic diversity of WSMV in our region and the possible impact on host resistance responses are unknown. We obtained two TriMV sequences, one complete genome from Kit Carson county (MT762125) and one partial sequence from Bent county. Both sequences had high identity (99%) with other available sequences, showing the low genotypic diversity among TriMV isolates.

**Fig. 4.**
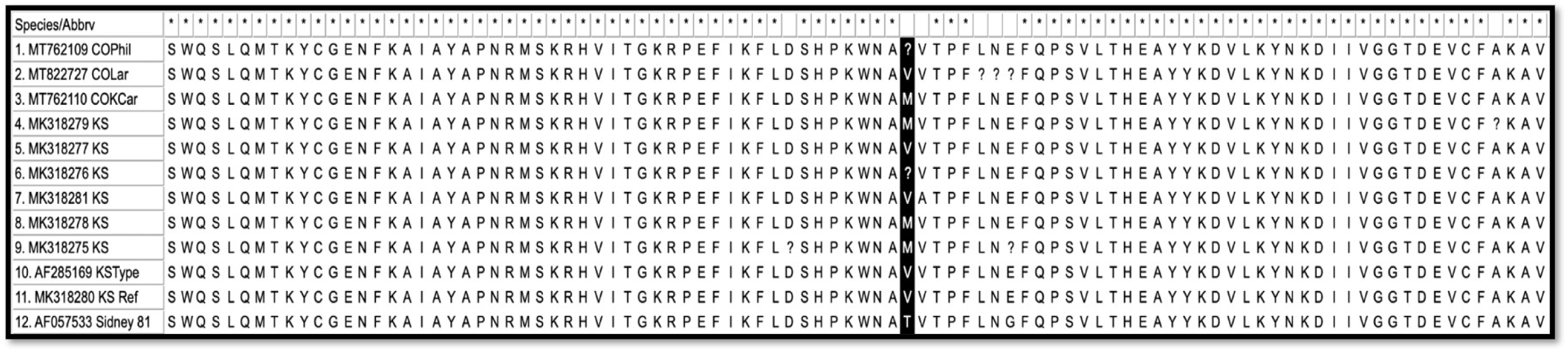
Amino acid variability in WSMV polyprotein amino acid sequences of Colorado isolates compared to seven Kansas isolates (Fellers et.al. 2019) and the reference strains, Sidney81 (AF057533) and Type (AF285169). Black box highlights common change between Sidney 81 and the other isolates reported in Fellers et.al. 2019. Alignments and visualization was performed using ClustalW.

HPWMoV was only detected in one location, Phillips county, which also harbored a diversity of other viruses (Table 3). Interestingly, we found 2-4 versions of all 8 segments of HPWMoV possibly composing at least two co-infecting emaravirus with complete genome segment complements. While the group with KS/NE/OH isolates (hereafter called group A) are known to have two variants of the RNA3 segment of HPWMoV (Stewart 2016), this is the first report of multiple variants for all eight RNA segments in a single sample. Fourteen of these segments contain complete coding sequences and have been deposited in GeneBank under accessions MT762111-MT762124. Phylogenetic analysis of the complete sequences of the nucleoprotein encoding RNA3 segments of members of both groups of HPWMoV and three variants all from the Phillips county sample, shows version Colorado RNA3C (MT762120) is similar to isolates from OH/TX or group A (Fig. 5). The other two variants are divergent from both groups of HPWMoV with Colorado RNA3A (MT762122) having only 75% identity with a member of group A and RNA3B (MT762121) having 74% identity with a member of group B; however Colorado RNA3A and RNA3B are similar to each other (Fig. 5). We also detected four variants of the RNA5 segment, three complete (MT762117-MT762115) and one partial sequence. These data suggest the presence of a new HPWMoV variant in Colorado in addition to a variant that is highly similar to known group A isolates.

**Fig. 5.**
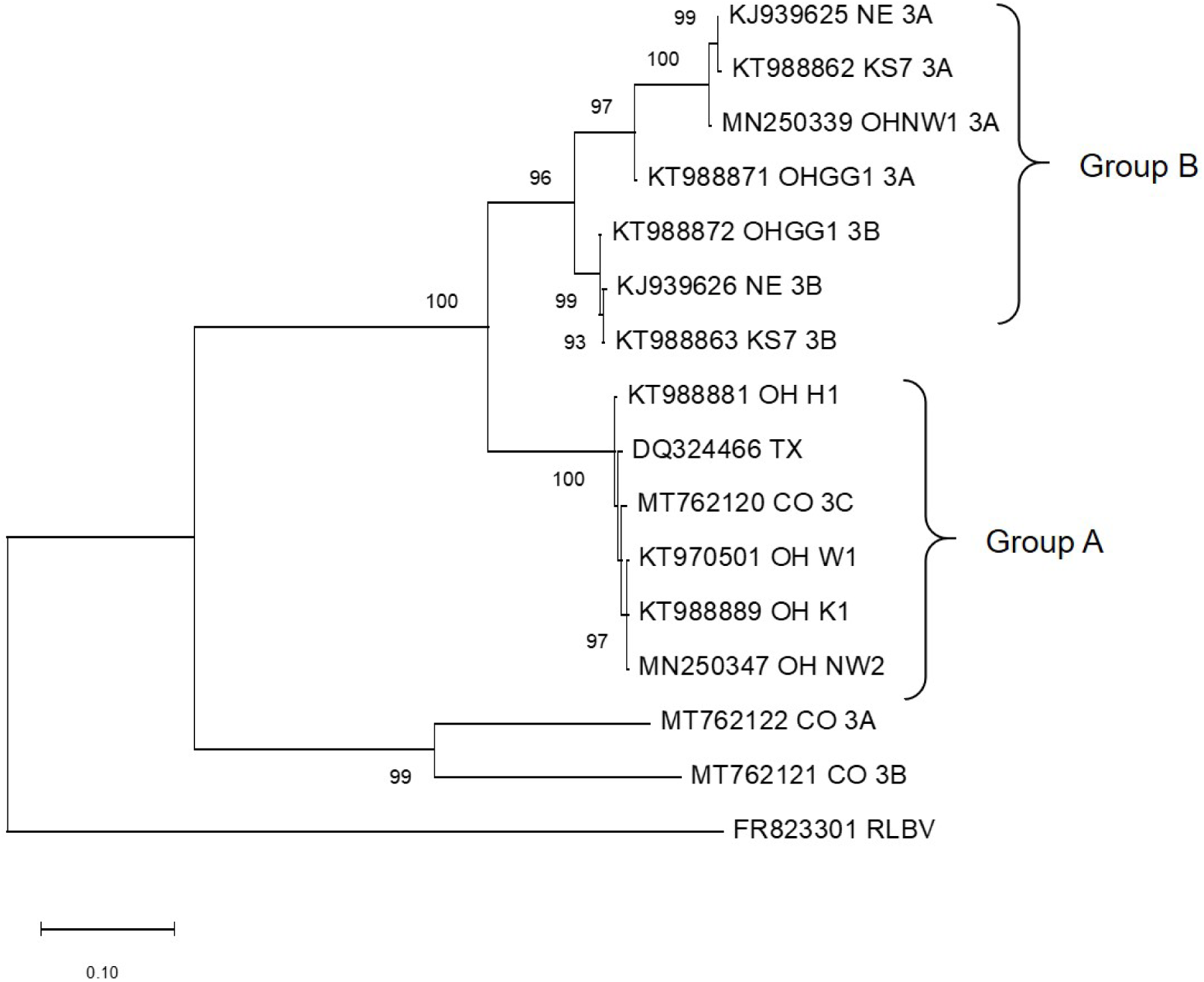
Phylogenetic tree of High Plains wheat mosaic virus (HPWMoV) isolates using the complete RNA3 segment encoding the nucleoprotein. Scale bar indicates percent genetic distances. Phylogenetic analysis by maximum likelihood method was based on a sequence alignment using ClustalW in MEGAX. Bootstrap values less than 70% out of 1000 replicates are not shown.

### Virus Resistance in Variety Trial

Wheat varieity trial included a combination of 24 public and private varieties and experimental lines. Seed companies with entries in the variety trials included AgriMaxx Wheat, AgriPro Syngenta, Dyna-Gro Seed, Limagrain Cereal Seeds, and WestBred Bayer. There were entries from the Colorado marketing organization, PlainsGold. The gerplasm included varieties with no known resistance, a single resistance marker to either WCM (*WCM6D*) or WSMV (*Wsm2*), and one variety, Guardian, with both resistance markers, *WCM6D* and *Wsm2* (Table 4). There were significant differences in virus titer (log copies of WSMV/mg of leaf sample) among varieties (F= 2.90; df=24,2; *P*=0.0007) (Fig. 6). The variety Guardian had lowest virus titer albeit not statistically different from others such as Snowmass 2.0, CO13D0346, CO15D098R that harbored a single resistance marker. Thunder CL and WB-Grainfield were statistically in the same group as the above four, but do not contain any resistance markers. While TriMV was detected in these samples, there was no significant differences between varities (F= 1.65; df=24,2; *P*=0.06).

**Table 4.** Presence of virus (*Wsm2*) and mite resistance (WCM 6D) genes in varieties tested at the Colorado State University Irrigated Variety Performance Trial. Blanks indicate that presence of resistance marker was not tested in these lines.

| Variety name | *Wsm2* | WCM 6D |
| --- | --- | --- |
| AM Eastwood | -- | -- |
| Brawl CL Plus | negative | negative |
| Breck | negative | negative |
| Canvas | negative | positive |
| CO13D0346 | positive | negative |
| CO15D098R | negative | positive |
| Crescent AX | negative | positive |
| Denali | negative | negative |
| Guardian | positive | positive |
| Long Branch | -- | -- |
| Monarch | negative | negative |
| Snowmass 2.0 | positive | negative |
| Sunshine | negative | negative |
| SY Wolf | -- | -- |
| SY Wolverine | -- | -- |
| Thunder CL | negative | negative |
| WB4269 | Negative | negative |
| WB4303 | negative | negative |
| WB4418 | -- | -- |
| WB4595 | -- | -- |
| WB4699 | -- | -- |
| WB-Grainfield | negative | negative |

**Fig. 6.**
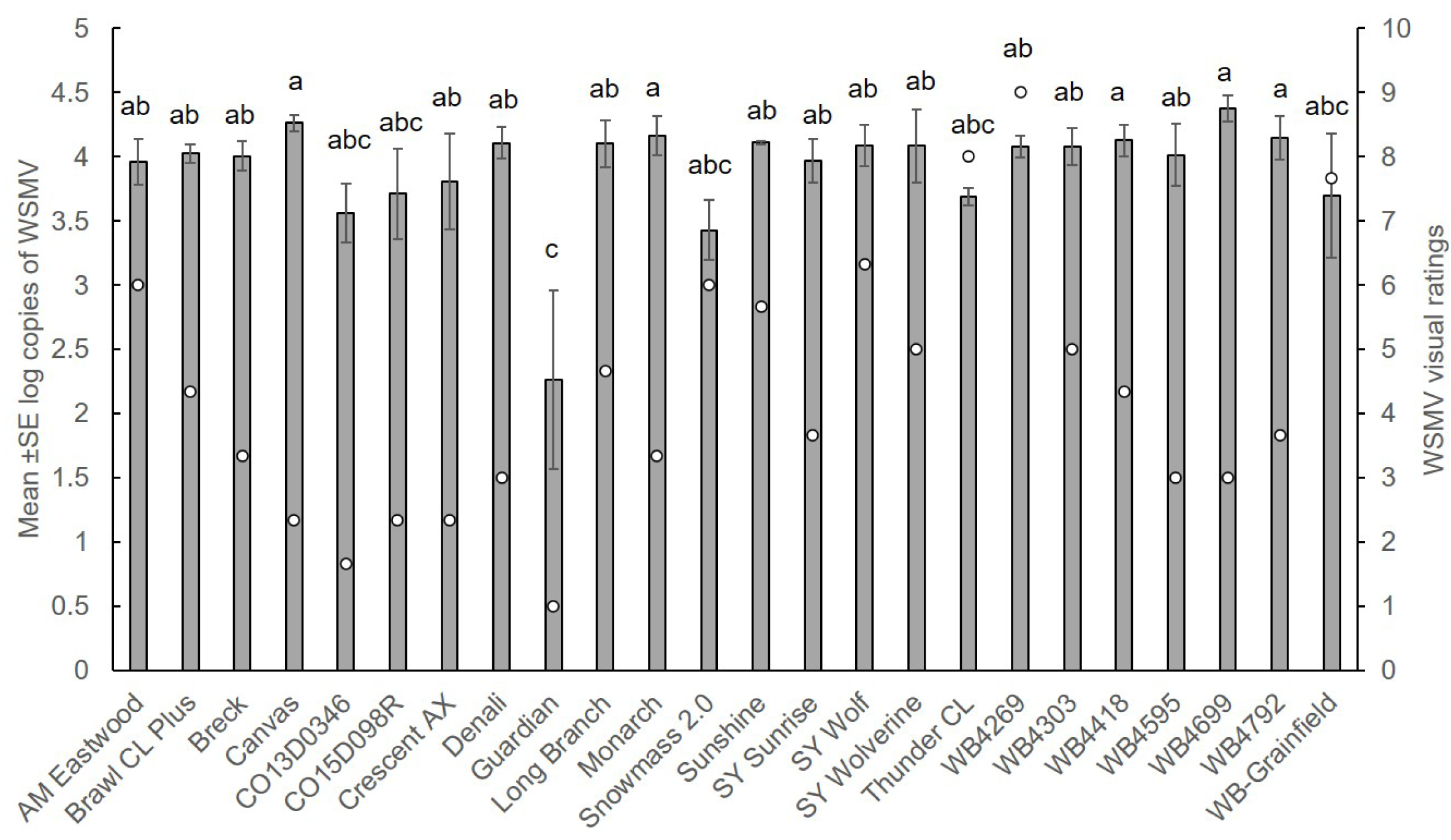
Response of wheat varieties and CSU advanced breeding lines to natural infection of wheat streak mosaic virus (WSMV) in an irrigated variety trial. Bars indicate mean of three biological replicates ± SE log copies of WSMV per variety. Circles indicate average WSMV visual rating on a 0-9 scale where 1= no damage and 9=severe damage. Different letters indicate significant differences between varieties at *P*<0.05.

## Discussion

Mite-vectored wheat viruses continue to cause significant yield losses in Colorado. To date, there has been no information on wheat curl mite-virus pathosystem in Colorado. Research is needed to increase our understanding of the biology, ecology and epidemiology of the WCM vector and three important mite-transmitted viruses, WSMV, TriMV and HPWMoV. Moreover, there is no information on emerging viruses in wheat, thus metagenomic sequencing can greatly enhance virus detection and characterization in wheat. In the current study, we identified the presence of two WCM genotypes, Type 1 and Type 2 from six populations collected from Colorado. We found rich sequence diversity of WSMV isolates and HPWMoV isolates collected from Colorado, whereas TriMV isolates had minimal sequence diversity. Wheat virome analysis confirmed the presence of known viruses such as WSMV, TriMV and HPWMoV, but also revealed presence of several mycoviruses and a novel Ixeridium yellow mottle virus 2 from Colorado wheat samples. Analysis of Colorado wheat germplasm showed that wheat varieites that contained both WCM and virus resistance genes had lower WSMV titer compared to varieties with only one resistance marker.

The variability in the WCM populations in a region can affect the prevalence and severity of virus infection (Wosula et al. 2016) and responses to WCM resistance genes (Dhakal et al. 2017; Harvey et al. 1999). The WCM is a cryptic species complex that includes two globally distributed lineages: Type 1 and Type 2 that are distinguishable using mitochondrial (mtDNA COI, 16S) and ribosomal (28S rDNA D2, ITS1–ITS2) marker and also differing in their host use patterns (generalization versus specialization) (Carew et al. 2009; Hein et al. 2012; Skoracka et al. 2014; Skoracka et al. 2018; Wosula et al. 2016). In the current study, we used the ITS1 sequence to understand phylogenetic relationships among mite populations from this study plus populations from within the U.S including KS, SD, MT, TX, NE, and outside the U.S, such as France, Australia, Argentina and Brazil. The WCM populations collected from Sedgwick and Larimer county 1 showed 100% sequence similarity as that of Type 1 mites from TX, NE, MT and SD. In contrast, mites collected from Larimer county 2 (same location as Larimer county 1 but at different time points), Kit Carson, Adam and Philips counties had ITS1 sequence identical to that of Type 2 mites from Nebraska (Fig. 1). The mitochondrial COI sequencing also demonstrated the two distinct types of WCM populations (Hein et al. 2012). These results suggest that both genotypes of mites are present in Colorado; moreover, mites from Larimer county, collected from the same area but during different times of the growing season belonged to both genotypes, which suggest that mixed populations can occur within fields similar to that observed for other wheat producing regions of the U.S. Great Plains (Siriwetwiwat 2006).

In the past, ELISA was the standard method for detection of mite-vectored viruses of wheat; however the more sensitive real-time qPCR analsysis is likely to increase the detection limit thereby modifying the data on virus incidence and prevalence in a given region (Bryan et al. 2019). In the current study, we used qPCR anslysis to detect WSMV and TriMV in wheat samples collected across Colorado. The most significant finding was that WSMV was detected in 95% of the virus-positive samples, followed by coinfection of WSMV and TriMV (19%) and WSMV and HPWMoV (8%). Single infection of WSMV was more frequent than coinfections, which is similar to previous findings (Burrows et al. 2009; Byamukama et al. 2013). Coinfections of WSMV with TriMV, and WSMV with HPWMoV occurred somewhat frequently, which is beneficial to know from the growers’ standpoint because WSMV and TriMV act synergistically in wheat resulting in more severe symptoms and yield losses compared to single infections (Byamukama et al. 2012; Tatineni et al. 2010). No single infection of TriMV and HPWMoV or coinfection of these viruses were detected.

The genetic diversity of WSMV has been evaluated among various isolates in the U.S. and from around the world by sequencing the coat protein (CP) (Robinson and Murray 2013) and more recent whole genome sequencing (Schubert et al. 2015). Based on the CP sequence, virus isolates have been divided into two clades, clade I and II. Isolates in clade I shared sequence similarity with isolates from Europe and isolates in clade II were similar to isolates originating from Australia, Argentina, and the American Pacific Northwest (Robinson and Murray 2013). Variability based on WGS revealed three clades, clade A, B and D (Schubert et al. 2015). Clade A represents isolates from Mexico, known as El Batán and clade B contains isolates from Europe, Russia and Iran. Clade D includes isolates from North and South America, Australia, Canada and Turkey. Phylogenetic analyses using a portion of the WSMV NIb region from Colorado isolates and WSMV whole genome sequences revealed significant diversity among Colorado isolates (MT465688-MT465692) within clade D; however, two isolates, MT465691 collected from Larimer county and MT762109 collected from Phillips county appeared to be genetically distinct from other isolates in the U.S. In addition, another isolate collected from Larimer county, MT465688 was similar to a new variant of WSMV (MK318278) collected from Kansas (Fellers et al. 2019). Because this isolate was collected from volunteer wheat on field margins, conclusions about potentially resistance-breaking nature of this isolate could not be made. Another isolate from Kit Carson county was collected from Snowmass 2.0 which contains the resistance gene, *Wsm2*. While genotypic factors likely play a role in host resistance response, other factors, such as temperature and coinfection with other viruses may also contribute to breakdown of the resistance observed in varieties containing *Wsm2*. Indeed, four isolates collected from *Wsm2*-harboring varieties from Kansas described in Fellers et al. (2019) also harbored multiple viruses including TriMV and/or Barley yellow dwarf virus (BYDV). Similarly, the isolate from Kit Carson county was coinfected with TriMV in the current study. Future research should be directed towards screening Colorado’s elite germplasm with these new variants to determine potential for resistance breakdown. Overall, these data indicate considerable diversity in WSMV isolates in our region, which could make breeding for durbale resistance difficult if there is differential response to WSMV resistance genes to different isolates.

To date, a handful of studies have detected viruses in wheat using NGS. A novel polerovirus named wheat leaf yellowing-associated virus (WLYaV) was identified from China (Zhang et al. 2017). Fellers et al. (2019) used Oxford Nanopore sequencing technology (ONT). to confirm the presence of important wheat viruses and to identify bacteriophages. More recently, Singh and colleagues reported the first record of two species of cereal yellow dwarf virus and wheat yellow dwarf virus (family *Luteoviridae/* genus *Polerovirus*) in wheat in the Czech Republic (Singh et al. 2020). In the current study, we identified 10 viruses in the wheat virome, including the three WCM-transmitted viruses, WSMV, TriMV and HPWMoV. This supports our previous findings based on qPCR analysis. Interestingly, 2-4 versions of all 8 HPWMoV segments were detected, which suggests that it is a co-infection of at least 2 emaraviruses. Multiple infections involving variable numbers of genome segments has been described for snake-infecting reptarenavirues, which, like the emaraviruses, belong to the *Bunyavirales* order (Stenglein et al. 2015). Two of the nucleoprotein encoding RNA3 segments were divergent from the two known groups of HPWMoV suggesting the presence of new variants of HPWMoV in Colorado. Future research is needed to understand the biological significance of the different groups of HPWMoV on host response. In addition, a novel virus, Ixeridium yellow mottle virus 2-like that is tentatively an umbravirus was identified.

Lastly, several mycoviruses or fungi-infecting viruses were identified including; *Plasmopara viticola* (the causal agent of grapevine downy mildew disease) associated viruses, Fusarium poae negative-stranded virus 2-like and Coniothyrium diplodiella negative-stranded RNA virus 1. These likely correspond to viruses infecting a *Fusarium* spp., which was present in this dataset as the most abundant non-host taxon identified. Morover, Fusarium poae negative-stranded virus 2 has been isolated from *Fusarium poae* strain SX63 (Wang et al. 2016). Our understanding of mycoviruses is poor relative to our understanding of plant viruses. Most mycoviruses do not cause any morphological changes in their fungal hosts (Ghabrial and Suzuki 2009; Wang et al. 2013). However, some mycoviruses such as Fusarium graminearum virus 1 can lead to devastating effects in their pathogenic fungal hosts including, reduced mycelial growth, decreased spores and/or sclerotia production, suppression of secondary metabolites, and attenuated virulence. This suggests mycoviruses could be a promising biocontrol agent for combating fungal diseases (Nuss 2005). The phenomenon by which mycoviruses reduce the ability of their fungal hosts to cause disease in plants is known as hypovirulence. (Dawe and Nuss 2013; Nuss 2005; Pearson et al. 2009). In contrast, a virus-infected fungus confered thermal tolerance to host plants, suggesting that mycoviruses could participate in mutualistic three-way symbioses (Márquez et al. 2007). Overall, our study provides a novel insight into the diversity of viral communitites including mycoviruses present in wheat. Additional sampling of mycovriuses could reveal novel candidates for biocontrol of plant pathogenic fungi. Future research may be aimed at understanding the diversity and dynamics of these viruses and mycovirus–host interactions.

Mite-vectored wheat viruses have been controlled by cultural practices and genetic resistance to the mite and pathogen (Tatineni and Hein 2018). In the current study, we screened a diverse germplasm including public and private varieties and experimental lines to natural infestation of WCM and mite-transmitted viruses. The genotype of the mites at the location of the variety trial was Type 2, which is the more virulent of the two genotypes. Our results demonstrate that varities with WCM or virus-resistant marker were effective in reducing WSMV levels. The WCM resistance was attributed to a novel gene mapped onto chromosome arm 6DS originally identitied in TAM112 (Dhakal et al. 2018). The resistance to WSMV was due to *Wsm2* gene identified by Haley et al. (2002) and is incorporated into several commercial varieties (Haley et al. 2011). With increasing acreage of varieities containing *Wsm2*, it is likely that isolates that can overcome resistance will be selected. A *Wsm2* breaking WSMV variant has been isolated from foxtail (Kumssa et al. 2019). There have been reports of increasing mite populations and virus infection in resistant varieties (Tatineni and Hein 2018). Indeed, we found variation in WSMV resistance among varieites that contained the mite and virus resistance markers. This may in part be due to heterologous effect of *Wsm2* (Chuang et al. 2017) in some lines or because the lines were derived from different genetic background. The only variety that harbored dual resistance markers (*WCM6D* and *Wsm2*), Guardian had lowest WSMV titer, albeit not statistically significant from varieites that harbored single resistance marker. This suggests that pyramiding mite and virus resistance genes can provide enhanced protection and improve durability. However, increased deployement of these resistant varities will likely cause the mite vector and virus to overcome these resistance mechanisms. This highlights the need for a multi-faceted approach to overcome the disease complex which includes managing alternate or “green bridge” hosts of mites, avoid early planting, planting resistant varieties, and continued search for novel sources of resistance.

## Acknowledgements

This research is supported by funds from Colorado Wheat Research Foundation and Colorado Wheat Administrative Committee. We would like to thank the extension agents and wheat growers for sending us samples.

